# Intermittent Cyclic Stretch of Engineered Ligaments Drives Hierarchical Collagen Fiber Maturation in a Dose- and Organizational-Dependent Manner

**DOI:** 10.1101/2024.04.06.588420

**Authors:** Leia D. Troop, Jennifer L. Puetzer

## Abstract

Hierarchical collagen fibers are the primary source of strength in tendons and ligaments, however these fibers do not regenerate after injury or with repair, resulting in limited treatment options. We previously developed a culture system that guides ACL fibroblasts to produce native-sized fibers and fascicles by 6 weeks. These constructs are promising ligament replacements, but further maturation is needed. Mechanical cues are critical for development *in vivo* and in engineered tissues; however, the effect on larger fiber and fascicle formation is largely unknown. Our objective was to investigate whether intermittent cyclic stretch, mimicking rapid muscle activity, drives further maturation in our system to create stronger engineered replacements and to explore whether cyclic loading has differential effects on cells at different degrees of collagen organization to better inform engineered tissue maturation protocols. Constructs were loaded with an established intermittent cyclic loading regime at 5 or 10% strain for up to 6 weeks and compared to static controls. Cyclic loading drove cells to increase hierarchical collagen organization, collagen crimp, and tissue mechanics, ultimately producing constructs that matched or exceeded immature ACL properties. Further, the effect of loading on cells varied depending on degree of organization. Specifically, 10% load drove early improvements in mechanics and composition, while 5% load was more beneficial later in culture, suggesting a cellular threshold response and a shift in mechanotransduction. This study provides new insight into how cyclic loading affects cell-driven hierarchical fiber formation and maturation, which will help to develop better rehabilitation protocols and engineer stronger replacements.

## 1. Introduction

Tendons and ligaments are critical for movement and stability, providing the strength for load transfer from muscle-to-bone and bone-to-bone, respectively. Tendons and ligaments are able to transfer and withstand these loads primarily due to hierarchically organized and aligned collagen fibers that run the length of the tissue [1–4]. Cells produce these hierarchical fibers by assembling tropocollagen molecules into fibrils (10–300 nm diameter), fibers (>10 μm diameter) and fascicles (100s μm to mm diameter) [5–7]. This hierarchical structure, along with collagen crosslinking and fiber crimp, provides tendons and ligaments with strong, non-linear, mechanical properties that are essential for long-term function [3,5,8,9]. The importance of these fibers to function are well understood, but replicating this hierarchical organization after injury or in engineered tissues remains a challenge [10,11].

Injuries to tendons and ligaments disrupt the collagen organization resulting in loss of function, pain, and decreased mobility [11,12]. Collagen fibers largely do not regenerate after injury or with repair, often resulting in unorganized scar tissue with reduced tissue function compared to healthy tissue [1,11]. There are more than 32 million tendon and ligament injuries per year in the United States [14] with limited repair options. In particular, the anterior cruciate ligament (ACL), which helps to stabilize the knee, is one of the most commonly injured ligaments resulting in an estimated 150,000 reconstructions each year in the US alone [15,16]. The current gold standards for ACL repair are autograft or allograft replacement. In addition to limited availability, these treatments have major drawbacks, including risk of donor site morbidity for autographs, immune response for allographs, and high risk of re-rupture [5,11,13,14,17–18]. Engineered replacements are promising, however, it remains a challenge to form the large hierarchically organized, crimped collagen fibers essential to long-term mechanical success [11,12].

Recently, we developed a novel culture system that guides ACL fibroblasts in unorganized high density collagen gels to develop aligned fibrils by 2 weeks of culture, which mature into native-sized hierarchically organized collagen fibers and early fascicles by 4 and 6 weeks [19,20]. These are some of the largest, most organized fibers produced to date *in vitro*; however, further maturation is needed for these constructs to serve as functional replacements.

Mechanical cues, including cyclic muscle activity, are critical for tissue development *in vivo* [1,5,21,22] and have been shown to drive maturation in engineered tissues *in vitro* [18,23–31]. In particular, intermittent cyclic loading at or below 5% strain is well established to improve fibril organization in engineered tendons and ligaments [1,18,22–30]; however the effect beyond the fibril level and on hierarchical fiber formation is largely unknown [1,10]. Further, the ACL has been reported to experience up to 12% strain in each gait cycle, suggesting strain rates above 5% may be beneficial for ACL fibroblasts, particularly beyond the fibril level which is less understood [1,11,32,33]. Additionally, there are large variations in optimal loading conditions for engineered tissues, most likely due to differences in scaffold material and design, which ultimately alter how the applied load is translated to cells [34]. Our culture system, which transitions from unorganized collagen at 0 weeks to aligned fibrils and fibers by 2 and 4 weeks, provides a means to explore how cells differentially respond to cyclic loading with different degrees of collagen organization. A better understanding of how cells respond to applied load at different levels of collagen organization could help to produce more uniform loading protocols across engineered systems and help to develop more optimal rehabilitation protocols.

The objective of this study was to investigate whether intermittent cyclic stretch, mimicking rapid muscle activity, could drive further maturation in our system. Additionally, we were interested in exploring whether 10% strain drives improvements over 5% strain in ACL fibroblast seeded constructs and whether cyclic loading had differential effects on cellular response when applied at different degrees of collagen organization. We hypothesize that intermittent cyclic loading will improve hierarchical collagen organization, composition, and tissue mechanics in a dose- and organizational-dependent manner, resulting in significantly stronger, functional ligament replacements.

## 2. Methods and Materials

### 2.1 Cell isolation

Ligament fibroblasts were isolated from neonatal bovine as previously described [12,20,35]. Briefly, neonatal (1-3 day old) bovine legs were obtained from a slaughterhouse within 48 hours of culling. The bovine cranial cruciate ligament (CCL), analogous to the human ACL and referred to as bovine ACL for the remainder of the manuscript, was aseptically isolated, diced, and digested for 15-18 hours in 0.2% collagenase. Cells were filtered, washed, counted, and frozen at 3 million cells/ml. Two separate isolations were performed, with each isolation having 3-4 donors pooled together. Prior to making constructs, cells were seeded at 2800 cells/cm^2^ and expanded 1 passage in Dulbecco’s Modified Eagle Medium (DMEM) media with 10% fetal bovine serum (FBS), 1% antibiotic/antimycotic, 50 µg/ml ascorbic acid, and 0.8 mM L-proline.

### 2.2 Construct Fabrication

High density, cell laden collagen gels were fabricated as previously described [12,20,35,36]. Briefly, type I collagen was extracted from an equal numbers of male and female Sprague-Dawley rat tail tendons (BIOIVT) and reconstituted at 30 mg/ml in a 0.1% acetic acid solution [12,37]. To generate constructs, collagen was mixed with phosphate buffer saline (PBS) and 1 N NaOH to initiate collagen gelation at pH 7 and 300 mOsm [38]. This solution was then immediately mixed with cells, ensuring cells were spread throughout the gel, cast into a 1.5 mm thick sheet gel, and allowed to set for 1 hour at 37 °C. The resulting sheet gels at 20 mg/ml collagen and 5 × 10^6^ cells/mL were cut into rectangles (8 x 30 mm) and divided between groups for culture. A different collagen stock was used for each sheet gel, with each sheet gel yielded 4-6 rectangular constructs. These constructs were distributed across experimental groups and time points. Thus, N refers to individual constructs produced from different collagen stocks and cell expansions.

### 2.3 Culture conditions and Mechanical Stimulation

One day after fabrication, static constructs were clamped into our culture device (**Fig. 1A**) as previously described [12,20], while loaded constructs were clamped into a modified CellScale tensile bioreactor (**Fig. 1B**), with both static and loaded constructs having a 20 mm gauge length between clamps. Loaded constructs were stimulated with an intermittent cyclic loading regime (**Fig. 1C**) established to generate matrix turnover and anabolic cellular response in engineered tissues [35,39–41]. Specifically, constructs were loaded with either 5% or 10% strain at 1 Hz for 1 hour on, 1 hour off, 1 hour on, 3x a week (Monday, Wednesday, Friday) to evaluate dose effect of cyclic loading (**Fig. 1C**). Culture media was the same as that used for cell expansion. Conditional media changes were performed immediately prior to loading by replacing half of the media during each media change throughout culture every 2-3 days [35].

**Figure 1:**
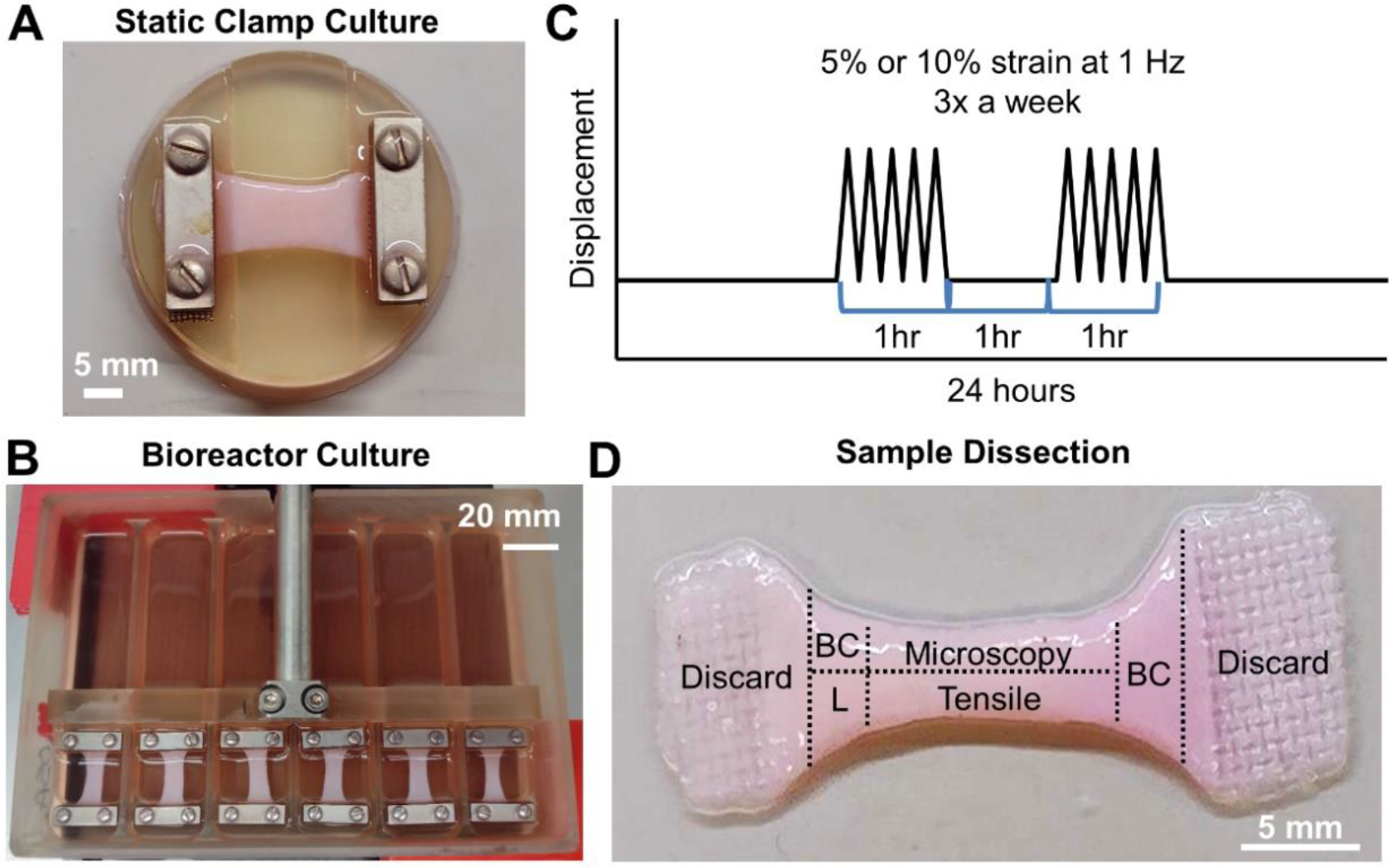
Experimental setup with constructs either statically clamped or cyclically loaded at 5 or 10% strain for up to 6 weeks. A) Static clamping culture device and B) bioreactor setup for cyclically loaded constructs. C) Depiction of intermittent loading regime. Constructs were loaded with an established loading regime, 3 times a week (MWF) for 1 hour on, 1 hour off, 1 hour on at 1 Hz and 5% or 10% strain. D) Depiction of tissue sectioning performed at each timepoint. Clamped regions were discarded and remaining tissue was allocated for biochemical composition analysis (BC), lysyl oxidase activity analysis (L), mechanical analysis, and microscopy analysis of organization.

### 2.4 Post-culture Analysis

Time points were taken at 0, 2, 4, and 6 weeks, with 0-week constructs taken out of culture 24 hours after 1 loading cycle. At each time point, 6-10 constructs per group were removed from culture, weighed, photographed, and sectioned for analysis of collagen hierarchical organization, composition, and mechanics as previously described (**Fig. 1D**) [21,25,39,42]. Due to significant contraction, not all sections for analysis could be obtained from loaded constructs, resulting in a greater number of constructs produced for loaded groups (N = 10) compared to control static constructs (N = 6). To track changes in contraction, construct percent weight and percent area were determined at each time point. Percent weight was calculated by comparing whole constructs wet weights to respective 0 week constructs. Percent area was determined by measuring construct surface area between the clamps in photographs taken at each time point using FIJI (NIH) and comparing to respective 0 week constructs, as previously described [20,35].

### 2.4 Post-culture Analysis

Time points were taken at 0, 2, 4, and 6 weeks, with 0-week constructs taken out of culture 24 hours after 1 loading cycle. At each time point, 6-10 constructs per group were removed from culture, weighed, photographed, and sectioned for analysis of collagen hierarchical organization, composition, and mechanics as previously described (**Fig. 1D**) [21,25,39,42]. Due to significant contraction, not all sections for analysis could be obtained from loaded constructs, resulting in a greater number of constructs produced for loaded groups (N = 10) compared to control static constructs (N = 6). To track changes in contraction, construct percent weight and percent area were determined at each time point. Percent weight was calculated by comparing whole constructs wet weights to respective 0 week constructs. Percent area was determined by measuring construct surface area between the clamps in photographs taken at each time point using FIJI (NIH) and comparing to respective 0 week constructs, as previously described [20,35].

### 2.5 Hierarchical Collagen Organization Analysis

Hierarchical collagen organization at the fibril (<1 µm), fiber (1-100 µm), and fascicle (>100 µm) length-scales were performed via scanning electron microscopy (SEM), confocal reflectance, and picrosirius red staining as previously reported [20,35,36]. Briefly, length-long sections of constructs and neonatal bovine ACLs were fixed in 10% formalin and stored in 70% ethanol. A total of 5-7 constructs per time point and condition and 5 neonatal ACL were analyzed via confocal reflectance. Following confocal analysis, a subset of 6 week constructs and neonatal ACLs were processed for SEM (N = 3) and picrosirius red staining (N = 3).

#### 2.5.1 Confocal Reflectance Imaging and Analysis

Confocal imaging for fiber level analysis was performed with a Zeiss LSM 980 microscope and Plan-Apochromat 20x/1.2 objective as previously described [12,31,39,43]. Briefly, a 405 nm laser was used to visualized collagen by capturing reflectance through a 27 µm pinhole at 400- 465 nm, while simultaneously a 488 nm laser was used to capture auto-fluorescence of cells at 509-571 nm. Five to eight representative 2D images were taken across the length of each construct and 3D images of 6 week constructs were obtained via Z-stacks with less than 1 µm step size and 18 - 22 um depth. Z-stacks were visualized with the FIJI 3D viewer plugin.

Confocal 2D images of constructs and native ACLs were analyzed using a custom Fast Fourier transform (FFT) based MATLAB code to determine degree of collagen alignment and fiber diameter as previously described [43]. Collagen alignment was scored using an alignment index, where 1 is unorganized and 4.5 is completely aligned [43]. Five to eight images per sample were analyzed and averaged to determine alignment and diameter for each construct. The reported alignment and diameter values are the average and standard error of construct values (N = 5-7 for constructs, N = 5 for neonatal ACL).

#### 2.5.2 SEM Imaging and Analysis

SEM analysis of fibril level ( <1 µm scale) organization was performed on 6 week constructs and neonatal bovine ACL with a Hitachi SU-70 FE-SEM as previously described [20,35,36]. Briefly, samples were dried via critical point drying and coated with 0.025-0.035 kAngstroms Platinum. Samples were imaged at a working distance of 10 mm, with 5 kV, at 10,000X and 50,000X magnification, with at least 6 images taken per sample. Images at 50,000X were analyzed to determine alignment and fibril diameter. Degree of alignment was measured by determining dispersion using the FIJI directionality function as previously described [20]. Dispersion values were obtained for 6-8 images per sample and averaged to determine average construct dispersion. The reported dispersion values are the average of construct values (N = 3 for constructs and native ACL). To determine fibril diameter a total of 20 fibrils per image were measured using FIJI (total 120 fibrils per construct) and pooled to determine average fibril diameter for each construct [36]. All fibril diameters from N = 3 samples (n = 360) were pooled for violin plots to visualize spread of fibril diameters at 6 weeks. For statistical analysis of fibril diameter, 120 fibril diameters per sample were averaged to compare construct averages at 6 weeks (N = 3 for constructs and native ACL).

#### 2.5.3 Polarized Picrosirius Red Imaging

Histological analysis was performed to observe fascicle level (> 100 µm scale) organization and crimp as previously described [12,20,35,36]. Briefly, fixed constructs (N = 3) and neonatal bovine ACL (N = 3) were embedded in paraffin, sectioned, and stained with Picrosirius red. Constructs were imaged with a Nikon Eclipse Ts2R inverted microscope and Nikon Pan Fluor 10x/.30 OFN25 Ph1 DLL objective in linear polarized light at 10 and 30x magnification.

### 2.6 Mechanical Analysis

Tensile tests were performed as previously described [36,43]. Briefly, samples (N = 6-8) were collected along the length of each construct, frozen for storage, thawed in PBS with EDTA-free protease inhibitor prior to testing, measured, and stretched to failure with a BOSE ElectroForce 3200 System equipped with a 250 g load cell. Samples were loaded at a strain rate of 0.75% /second, assuming quasi-static load and ensuring failure between the grips. Tensile properties were determined via a custom linear regression-based MATLAB code [35]. Briefly, the toe and elastic moduli were determined by fitting a linear regression to the stress-strain curve in each region, ensuring an r^2^ > 0.999, and the transition point was defined as where these two linear regressions intersected. The maximum point on the stress-strain curve was used to determine the ultimate tensile strength (UTS) and strain at failure.

### 2.7 Compositional Analysis

Compositional analysis for DNA, glycosaminoglycan (GAG), and collagen content were performed as previously described [12,20,35,36]. Briefly, two sections per construct were taken for compositional analysis (**Fig 1D**) and the values were averaged to determine composition for each construct. Construct sections and neonatal bovine ACL samples were weighed wet (WW), frozen, lyophilized, weighed dry (DW) and digested in 1.25 mg/ml papain solution at 60 °C for 15- 16 hours (N = 6-8). DNA, GAG, and collagen were determined via a modified Quant-iT PicoGreen dsDNA assay kit (Invitrogen), 1,9-dimethylmethylene blue (DMMB) assay at pH 1.5 [44], and a modified hydroxyproline (hypro) assay [45]. Constructs from each experimental group retained similar percent wet weight throughout culture, so DNA, GAG, and hydroxyproline are reported normalized to sample wet weight.

To determine LOX activity in constructs, separate sections from constructs (**Fig. 1D,** N = 5-6) were placed in a 6 M Urea 10mM Tris-HCl solution at pH 7.4 with 1% protease inhibitor at each time point and frozen at -80 °C. A fluorometric LOX activity assay (Abcam, ab112139) was used to determine arbitrary units (A.U.) of LOX activity. The assay was run according to manufactures protocol with recombinant LOXL2 (R&D Systems) used as a positive control and results normalized to sample DNA determined via the Quant-iT Picogreen dsDNA assay.

### 2.8 Acellular Constructs

To determine if load alone, rather than cellular response to load, drives collagen organization, acellular constructs were cultured in the same manner as cell-seeded constructs. Acellular constructs were produced as described above, however, instead of cell-seeded media being added to the collagen mixture, media alone was mixed into the collagen solution to produce 20 mg/ml constructs. Acellular constructs were cultured statically clamped, or with 5% or 10% intermittent cyclic loading for up to 6 weeks as described above. At 0 and 6 weeks constructs were removed from culture (N = 4-6) and sectioned the same as cell-seeded constructs for analysis of collagen organization, mechanics, and composition (**Fig. 1D**). Specifically, collagen organization was assessed by confocal reflectance, mechanics were assessed via tensile tests to failure, and collagen concentration was assessed as a measure of composition, all done according to methods above for cell-seeded constructs.

### 2.9 Statistics

SPSS was used to confirm normality of data within each group. Following confirmation of normality, data were analyzed via 2-way ANOVA and Tukey’s post-hoc analysis with *p* < 0.05 as significant (SigmaPlot 14). Since fibril dispersion and diameter data were only collected at 6 weeks, this data was analyzed via 1- way ANOVA. All data are expressed as mean ± standard error (S.E.M.).

## 3. Results

### 3.1 Gross Morphology

Gross inspection revealed all constructs contracted with time in culture, with both 5 and 10% load constructs appearing to have accelerated and increased contraction compared to static constructs by 4 weeks (**Fig. 2A**). Percent area measurements confirmed this observation, with loaded constructs having a significantly lower percent area compared to static controls starting by 2 weeks (**Fig. 2B**). However, interestingly, percent weight measurements, based on the wet weight of constructs, revealed all constructs contracted similarly with time in culture to 30-40% their original weight (**Fig. 2C**). Loading did not significantly affect percent weight compared to static constructs, suggesting that while there were differences in surface area reduction, loaded and static samples retained similar mass.

**Figure 2:**
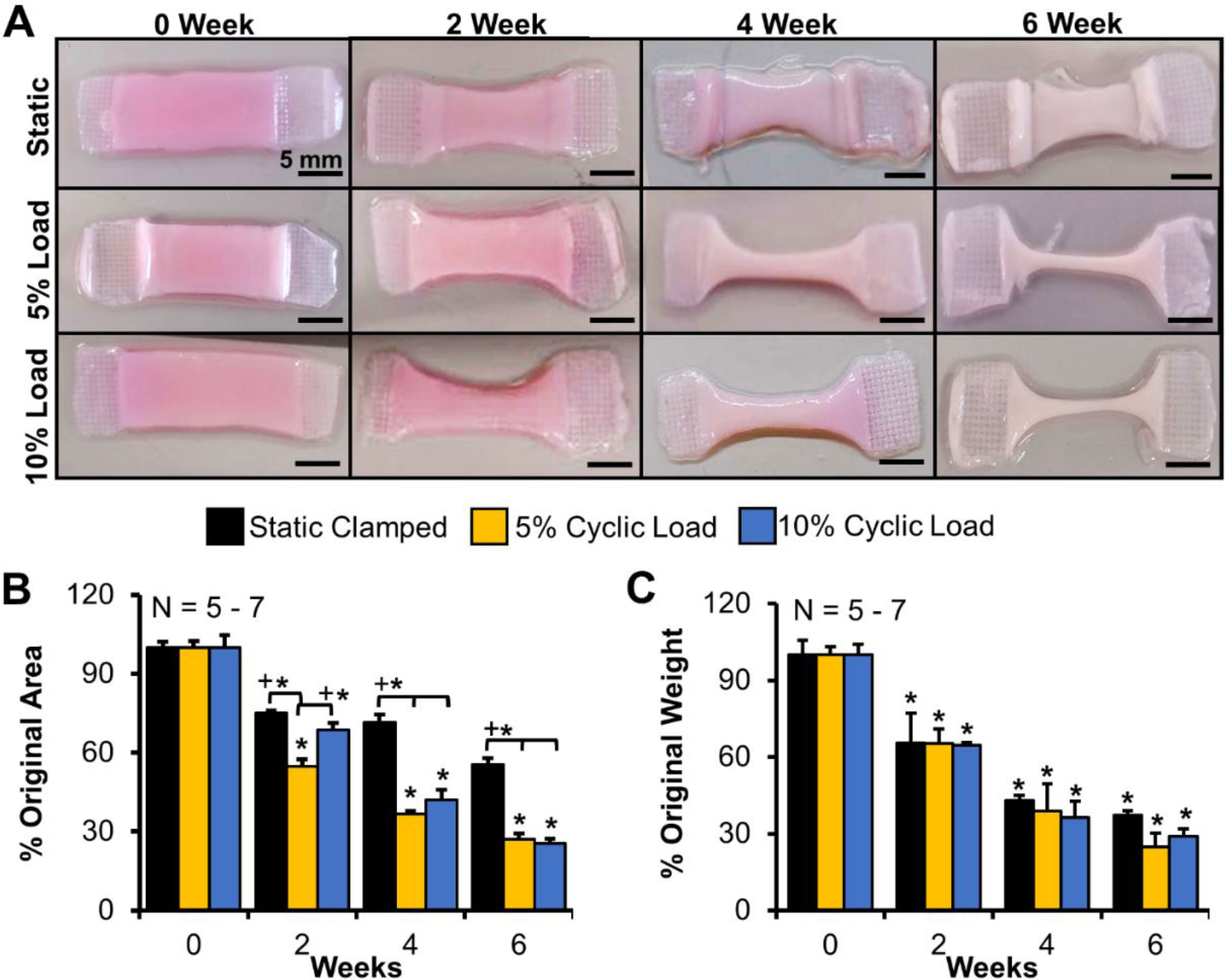
Loading appeared to increase construct contraction, but construct wet weight remained similar to controls. A) Photographs of representative constructs at each time point, B) percent area and C) percentage wet weight throughout culture compared to respective 0 week constructs. Scale bar = 5 mm, Data shown as mean ± S.E.M., Significance compared to *0 week or +bracket group (p <0.05).

### 3.2 Hierarchical Collagen Organization

Confocal reflectance analysis revealed all groups guided cells in unorganized collagen at 0 weeks to produce aligned collagen fibrils by 2 weeks and larger fibers by 4 and 6 weeks, similar to previous studies (**Fig. 3A**) [12,20,36]. Loading accelerated fiber development, with collagen fibers in both 5 and 10% constructs appearing larger and more organized by 4 and 6 weeks, similar to neonatal bovine ACL. In addition to organization, loaded constructs appeared to have developed crimp by 2 weeks, which appeared more uniform and regularly spaced by 6 weeks (**Fig. 3A, arrows**). Three-dimensional reconstructions of 6 week constructs further confirmed loaded constructs developed more clearly defined collagen fibers and increased crimp compared to static constructs, with 10% load constructs appearing to have larger, more organized fascicle formations compared to 5% (**Fig. 3A, Z-stacks**).

**Figure 3:**
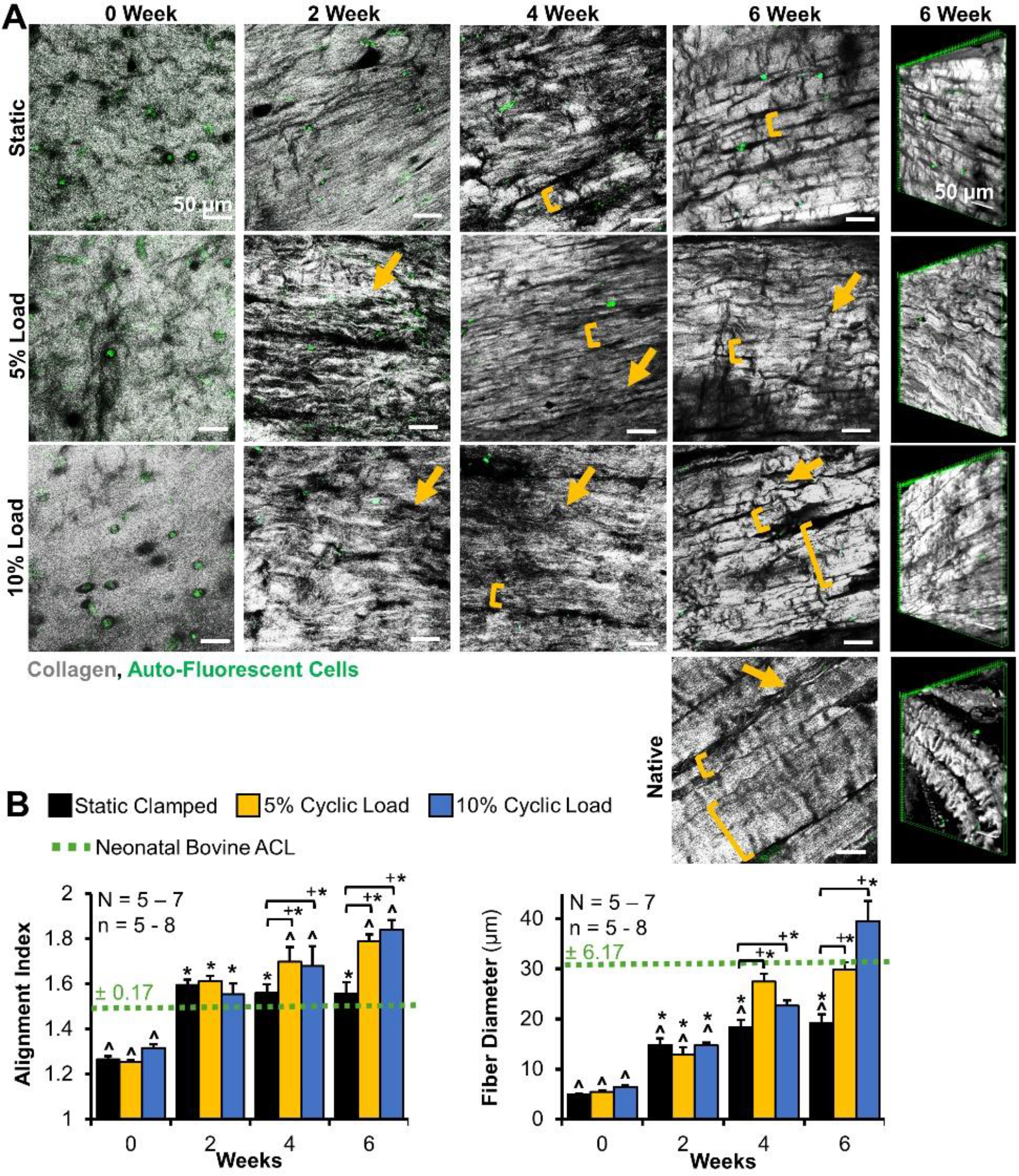
Loading increased fiber development in a dose-dependent fashion. A) Confocal reflectance revealed constructs develop aligned fibrils at 2 weeks, and larger fibers by 4 weeks (brackets), with loading producing increased fiber and fascicle formation (larger brackets) and crimp formation (arrows). Grey = collagen, green = cellular auto-fluorescence. scale bar = 50 μm. B) Degree of collagen alignment (reported via alignment index where 1 is unorganized, 4.5 is perfectly aligned) and average collagen fiber diameter determined via a FFT based image analysis. 5-8 images per construct were averaged to determine construct values, with 5-7 constructs analyzed per time point and 5 native tissue analyzed. Data shown as mean ± S.E.M., Significant difference compared to *0 week, ^native, or +bracket group (p <0.05).

Analysis of confocal images revealed that all groups had significantly improved alignment by 2 weeks, matching neonatal bovine ACL alignment (**Fig. 3B**). Loading further significantly improved collagen alignment over static controls and native tissue by 4 and 6 weeks. Average collagen fiber diameter also significantly improved for all groups with time in culture. Loading further improved fiber diameter in a dose-dependent manner. Both 5 and 10% load produced significantly increased fiber diameters compared to static constructs by 4 weeks, which were no longer significantly different from neonatal diameters. By 6 weeks, 10% load constructs had significantly larger fibers compared to 5% load and static constructs, reaching an average diameter of 38.9 ± 2.8 µm (**Fig. 3B**).

Similar to fiber level organization, SEM analysis revealed increased organization at the fibril level (<1 µm length-scale) in loaded constructs by 6 weeks (**Fig. 4**). Both 5 and 10% load appeared to drive increased alignment of fibrils compared to static controls, as well as compaction of fibrils into larger bundles similar to neonatal tissue (**Fig. 4A**). Image analysis confirmed that by 6 weeks, 5% and 10% load constructs had significantly decreased degrees of dispersion (i.e. increased alignment) compared to static controls (**Fig. 4B**). Further, measurements of fibril diameter revealed loaded constructs had significantly larger fibril diameters than static controls at 6 weeks, with 10% load constructs reaching neonatal ACL diameters (∼58 nm) and having significantly larger fibrils compared to 5% load constructs. Further, both loaded groups became more heterogeneous with a wide range of fibril sizes, similar to native tissue.

**Figure 4:**
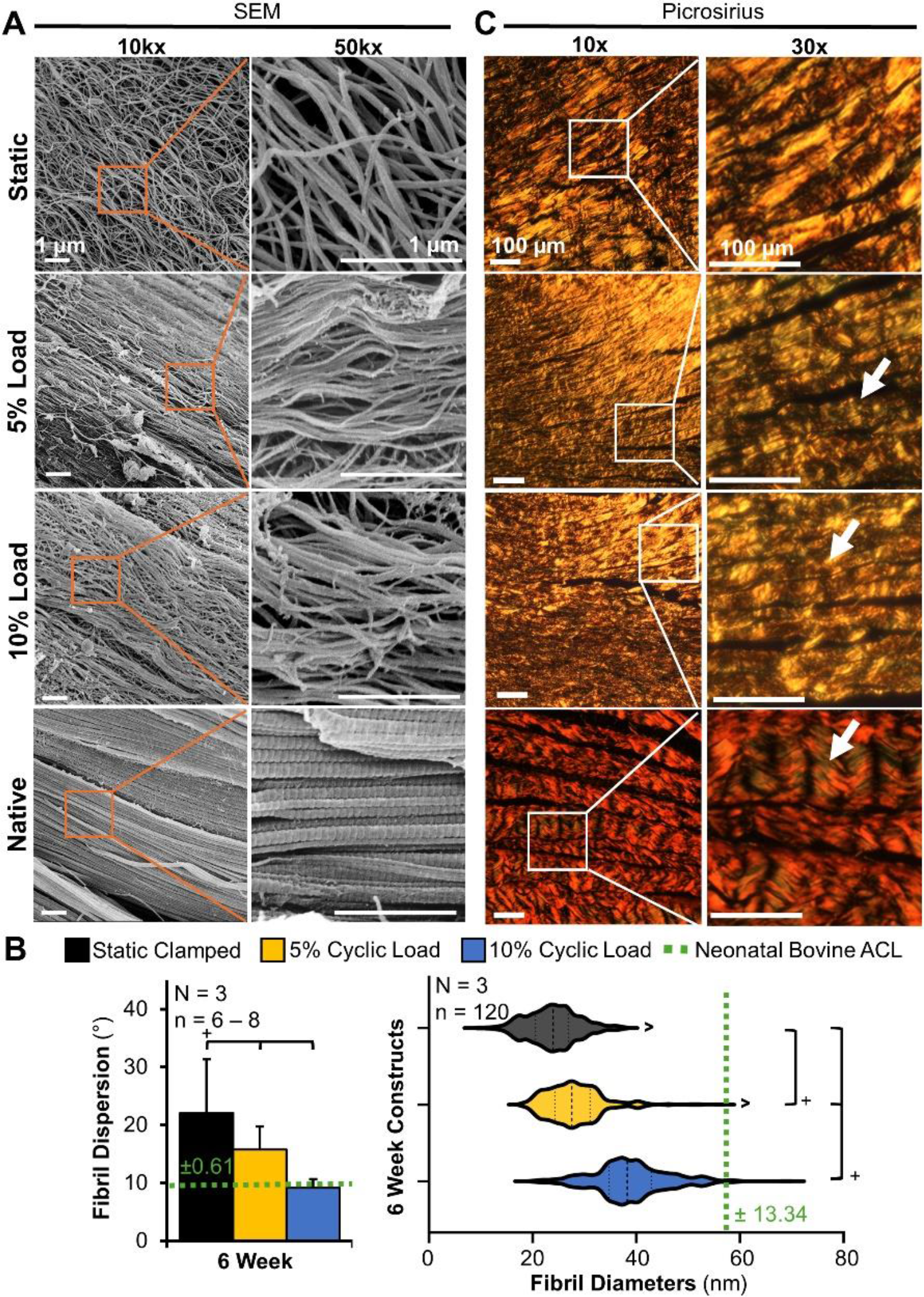
Loading increased fibril and fascicle formation in a dose-dependent fashion. A) SEM images of 6-week constructs to assess fibril length-scale organization (scale bar = 1 µm), and B) analysis of SEM images to determine fibril dispersion (lower dispersion indicating increased alignment), and fibril diameter of 6 week constructs. For dispersion, 6-8 images per construct were averaged to determine construct average (N=3 for constructs and native ACL). For fibril diameters, 20 fibrils per image were measured (total 120 fibrils per construct) and pooled to determine average fibril diameter for each construct. All fibril diameters from N=3 constructs were pooled (n = 360) to visualize spread of fibril diameters at 6 weeks. Data shown as mean ± S.E.M., Significance compared to ^native tissue or +bracket group (p <0.05) C) Fascicle length-scale organization at 6 weeks evaluated by picrosirius red staining imaged with polarized light (N = 3, scale bar = 100 µm). Loading contributed to enhanced fascicle formation and crimp formation (arrows).

Polarized picrosirius red analysis of the fascicle length-scale (>100 µm length-scale) further confirmed loading drove improvements in hierarchical collagen organization (**Fig. 4C**). Low magnification imaging (10x images) demonstrated loaded constructs had larger, more compact collagen bundles, indicative of early fascicle formation, compared to static controls. Higher levels magnification (30x images) revealed 5% and 10% load constructs had increased crimp formation compared to static controls, with 10% load constructs producing more pronounced, regularly spaced crimps than that of 5%. The regularity of crimp in loaded constructs approached the regularity and size of neonatal bovine ACL. Collectively, intermittent cyclic loading drove accelerated and increased hierarchical fiber formation, with 10% load producing regular crimps as well as increased fibril and fiber diameters that matched or exceeded neonatal bovine ACL.

### 3.3 Tissue Mechanical Properties

Mirroring improvements in organization, all constructs had significant improvements in tensile properties with time in culture (**Fig. 5**). Interestingly, despite 10% load constructs having improved hierarchical fiber formation, only 5% load constructs had improved elastic tensile properties over static controls by 6 weeks, with a significant increase in elastic modulus over static and 10% load groups at 6 weeks (**Fig. 5A**). Ultimately 5% load constructs had an elastic modulus of 3.5 ± 0.72 MPa at 6 weeks, surpassing reported immature 1 week old bovine ACL values (1-3 MPa [46]). Similarly, all constructs had improved UTS by 6 weeks, with 5% load constructs having a 2-fold greater UTS compared to 10% constructs by 6 weeks. Both 5% and 10% groups had decreased strain at failure compared to static samples by 6 weeks, suggesting increased maturation.

**Figure 5:**
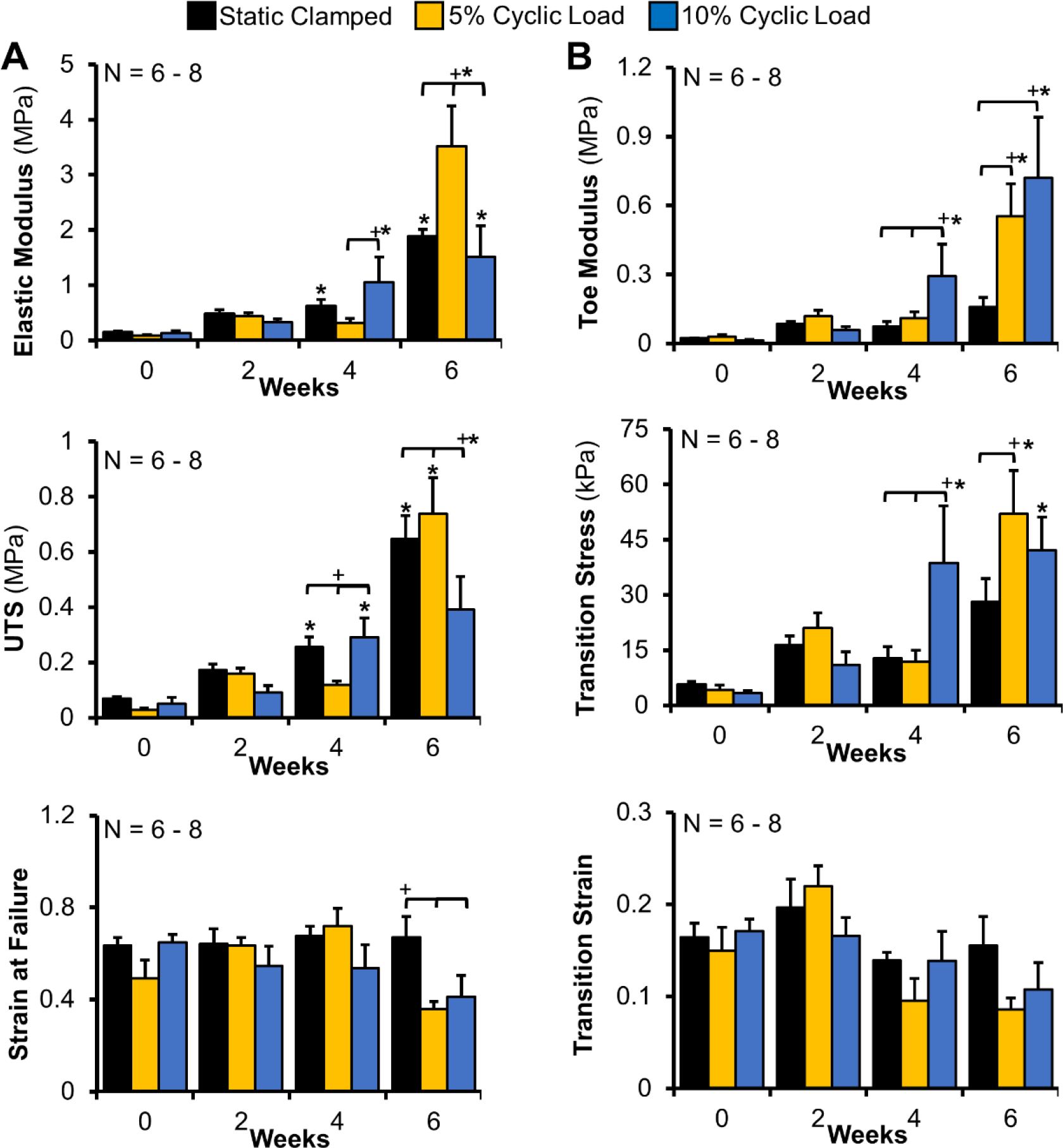
Loading increased mechanical properties in a dose-dependent fashion. A) 5% load constructs significantly improved elastic properties (Elastic modulus, ultimate tensile strength (UTS), and strain at failure) by 6 weeks compared to 10% load and static samples. B) 5% and 10% load constructs significantly improved toe-region properties (Toe modulus, transition stress, and transition strain) by 6 weeks, suggesting increased fiber alignment and development of functional crimp. N = 6-8. Data shown as mean ± S.E.M., Significance compared to *0 week and +bracket group (p < 0.05)

Despite 5% load only producing significant increases in elastic properties over static controls, both 5% and 10% load constructs had significant improvements in toe region properties (**Fig. 5B**), with 5% and 10% groups having a 2.5- and 3.5-fold improvement in toe modulus over static constructs, respectively. Further, 10% load constructs had accelerated improvements, with significant increases in toe modulus and transition stress by 4 weeks compared to static and 5% load constructs, but no significant change in transition strain.

### 3.4 Matrix Composition and LOX activity

With time in culture, all constructs had a significant increase in DNA, GAG, and collagen concentration (normalized to wet weight and dry weight), with the effect of loading on composition varying with time in culture and degree of collagen organization (**Fig. 6, Supplemental Fig. 1** for dry weight normalizations). Despite all groups having similar decreases in percent wet weight (**Fig. 2C**), loading appeared to increase cell proliferation resulting in significant increases in DNA normalized to wet weight at 2 and 4 weeks for both 5 and 10% load compared to controls, but this effect was lost by 6 weeks when all groups reach neonatal bovine ACL concentrations (**Fig. 6**). Interestingly, both load groups had a significant increase in GAG concentration at 0 weeks after just one loading cyclic, compared to static samples, and 10% load constructs had a further significant increase in GAG accumulation at 2 and 4 weeks (**Fig. 6**). However, by 6 weeks, all treatment groups had similar levels of GAGs, with both load groups leveling off at neonatal ACL concentrations. Collagen content represented by hydroxyproline, decreased in 10% load samples compared to static controls after one loading cycle and decreased in 5% load samples at 2 weeks compared to static controls, indicating that the cells may have initially been breaking down collagen in response to load. By 4 weeks, this trend was reversed, with both load groups increasing collagen concentration to static control levels, with 5% cyclic load constructs accumulated 2-fold more collagen than 10% load constructs. By 6 weeks, 5% load constructs reached neonatal bovine ACL collagen concentrations and had a significant 2-fold increase over static and 10% load groups (**Fig 6, Supplemental Fig 1C**). LOX activity was significantly increased by both 5 and 10% load by 6 weeks (**Fig. 6**). Interestingly, LOX activity only increased for 10% load constructs at 0 weeks after one loading cycle in unorganized collagen gels, and after 6 weeks of culture once aligned fibers had formed. Further, while 5% load significantly increased LOX activity at 4 and 6 weeks, 10% load constructs had a 2-fold increase over 5% load constructs by 6 weeks.

**Figure 6:**
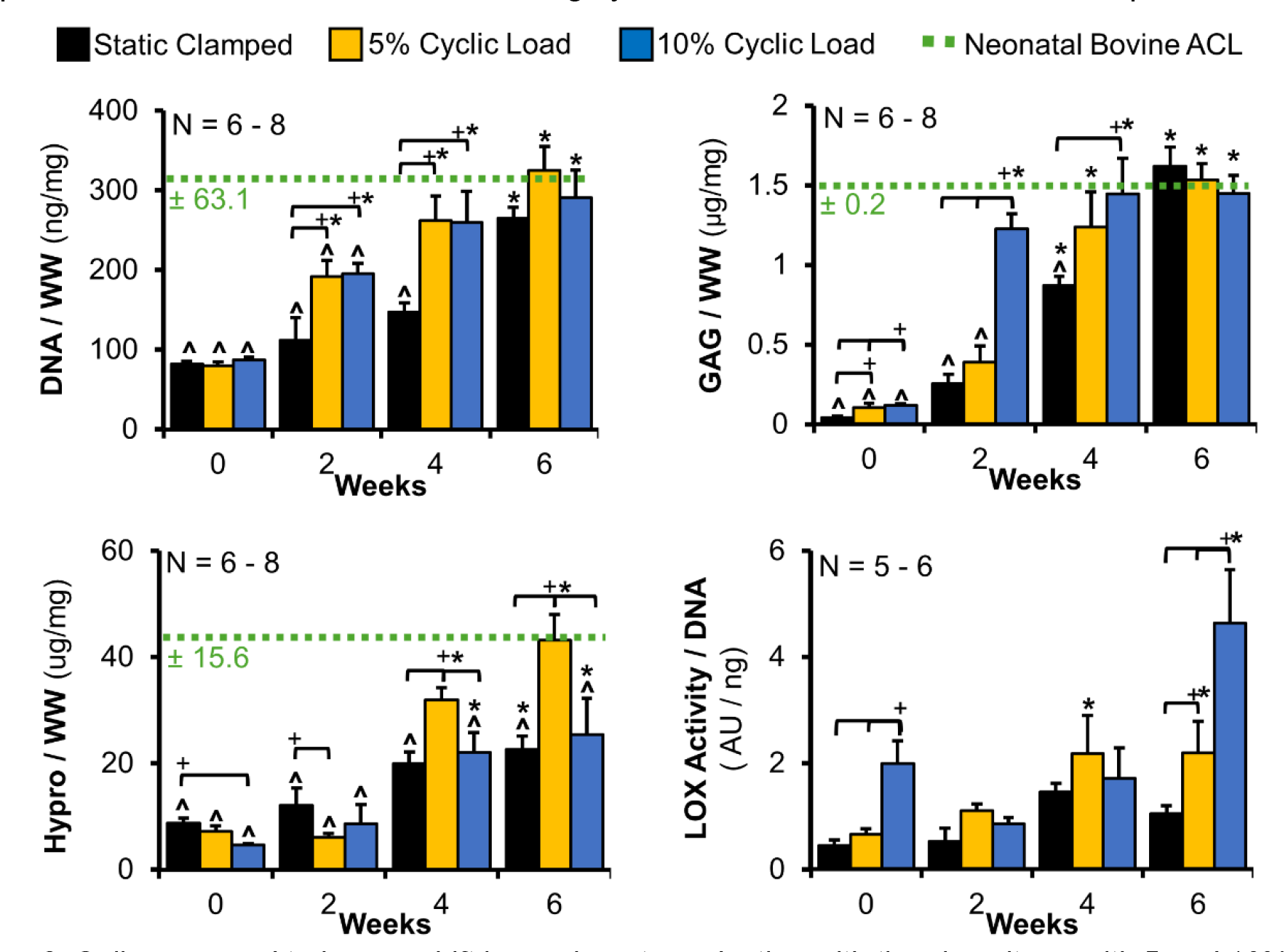
Cells appeared to have a shift in mechanotransduction with time in culture, with 5 and 10% load leading to differential changes in composition throughout culture. DNA, GAG, and collagen content, represented by hydroxproline, normalized to wet weight (WW, N = 6-8), and LOX activity in constructs normalized to DNA (N = 5-6). 5% load constructs matching neonatal ACL DNA, GAG, and collagen concentrations by 6 week. Data shown as mean ± S.E.M., Significance compared to *0 week, ^native, and +bracket group (p < 0.05).

### 3.5 Acellular Constructs

Acellular constructs were also cultured for up to 6 weeks to determine if stretching of collagen gels alone, rather than cellular response to load, drives construct maturation. All acellular constructs had no contraction and no change in percent weight by 6 weeks (**Fig. 7A**). Likewise, static controls and loaded acellular constructs had no fibril alignment or larger fiber organization by 6 weeks (**Fig. 7B**). Constructs remained unorganized throughout culture. Further, there was no accumulation or significant loss of collagen in any treatment group (**Fig. 7C**) and no improvements in tissue tensile mechanics (**Fig. 7D** and **Supplemental Figure 2**). Collectively, this data suggests that any improvements in organization and mechanics were due to cellular response to load and not induced organization from stretching of the collagen gels alone.

**Figure 7:**
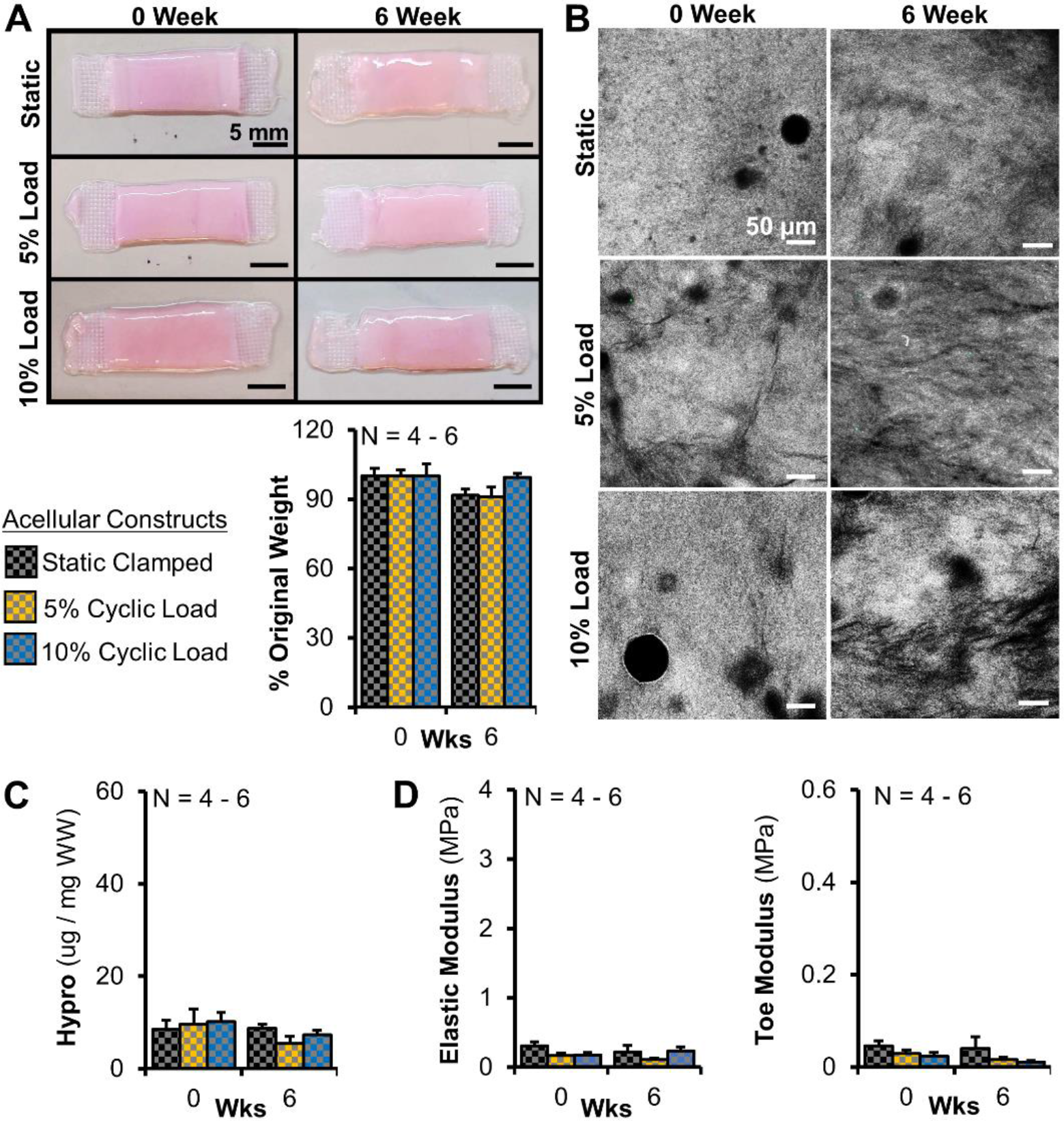
Loading had little-to-no effect on acellular constructs, demonstrating changes in cell-seeded constructs in response to load are cell-driven. Acellular constructs had no significant change in A) gross morphology and percent weight by 6 weeks of culture. B) Confocal reflectance at 0 and 6 weeks revealed little-to-no change in organization (Grey = collagen, scale bar = 50 µm). C) Collagen content, represented by hydroxproline and normalized to wet weight (WW), and D) Tensile properties (elastic and toe modulus) had no increases with time in culture (y-axis set to be similar scale used for cell-seeded constructs). Data shown as mean ± S.E.M., Significance compared to *0 week and +bracket group (p < 0.05).

## 4. Discussion

Collectively, this study demonstrates intermittent cyclic loading increases hierarchical collagen organization, crimp formation, and tissue mechanics. Both 5 and 10% load accelerated and increased hierarchical collagen fiber formation in a dose-dependent fashion, producing fibrils and fibers that matched neonatal bovine diameters by 6 weeks. Further, both magnitudes of load drove increased collagen crimp formation and increased toe-region mechanics, suggesting the development of crimp is leading to more functional mechanics. By comparing to acellular gels, we demonstrated these improvements in collagen organization and mechanics are cell driven, with the effect of cyclic load on cells varying depending on the degree of organization and magnitude of strain. Collectively, with respect to collagen production, cyclic load produced a more catabolic response early in culture while cells were in unorganized gels, and a more anabolic effect once cells were anchored on aligned fibrils, suggesting a shift in mechanotransduction with increased collagen organization. Further, while 10% load drove further improvements in fiber organization, only 5% cyclic strain produced increased elastic mechanics and collagen accumulation, ultimately matching neonatal ACL properties, suggesting a mechanical threshold in cellular response.

It is well established mechanical cues are critical for tissue development *in vivo* [1,5,21,22] and drive maturation of engineered tissues *in vitro* [18,23–31]. In particular, intermittent cyclic loading at or below 5% strain is well established to improve fibril alignment, fibril diameter, and collagen production in engineered tendons and ligaments [1,5,17,21–31,47]; however the effect at the fiber and fascicle length-scale is largely unknown [1,19,23,48,49]. In this study we found that intermittent cyclic loading accelerates and improves hierarchical collagen organization at the fibril, fiber, and fascicle length-scale (**Fig 3-4**). At the fibril level, similar to previous studies [47], 5 and 10% cyclic load led to increased fibril alignment and fibril diameter, with both matching neonatal ACL alignment and 10% load constructs matching neonatal native diameter by 6 weeks (**Fig. 4B**). Additionally, fibril diameters in both 5% and 10% load constructs became more heterogeneous by 6 weeks, similar to the shift observed in native fibril diameters during development [50]. At the fiber level, loading increased fiber alignment and diameter, with loaded constructs matching and exceeding neonatal ACL alignment by 2 weeks and fiber diameters by 4 weeks. At the fascicle length-scale, loaded constructs appeared to have improved fascicle formation with increased crimp formation by 6 weeks. Collectively, intermittent cyclic loading drove significant improvements at each hierarchical level over static controls, producing some of the largest and most hierarchically organized collagen fibers to date *in vitro*.

Further, we found that 10% cyclic loading stimulated cells to produce larger and more organized hierarchical collagen fibers in comparison to 5% load, with 10% constructs having increased alignment and diameters at the fibril and fiber level (**Fig. 3B & 4B**). Traditionally strains at 5% or below are reported to be optimal for tendon and ligament engineering [18,26–30], as most native tendons and ligaments experience strains under 5% [51]. However, interestingly, here we found that 10% cyclic strain led to improvements in fibril, fiber, and fascicle level organization compared to both static and 5% load constructs. This may be due to the ACL receiving larger strain *in vivo* (up to 10-12%) [33].

In addition to driving increased hierarchical organization, intermittent cyclic loading drove cells to produce collagen crimp starting at 2 weeks, with crimp appearing to increase in number and frequency by 6 weeks (**Fig. 3-4**). Crimps are wavy, planar zig zag structures in collagen fibers that serve a protective role in ligaments by taking on initial loads when strained [5,12,52,53]. Crimp formation is not well understood [7,54,55], but previous work has typically generated crimp structures by either manufacturing crimp into the scaffold prior to cell seeding [52,56–61], or by releasing restrained constructs post-seeding to allow for scaffold contraction [55]. Here, loading stimulated cellular development of crimp, evident by the fact that acellular constructs had no development of crimp even after 6 weeks of loading. *In vivo* crimp develops postnatally as cells gather fibers into larger bundles [62]. Specifically, it has been reported crimp increases with increasing strain and maturation *in vivo* [54,55,63,64]. Similarly, in this study we found increased crimp formation with applied dynamic cyclic strain and increasing hierarchical fiber maturation. It is not well understood how crimp forms, but it is thought to be through mechanotransduction and the mechanobiological environment [54,55]. It has been reported that collagen crimp in rat tail and Achilles tendon are fully extended at 4% strain *in vivo* [53,65]. Previously, studies applying strains <5% in scaffolds without pre-fabricated crimp have not reported any development of crimp [24,26,27], thus a strain greater than the natural crimp extension length of 4% may be needed to stimulate cells to produce crimp formation. This study demonstrates that application of intermittent cyclic loading is capable of driving cells in collagen gels to produce crimps approaching native structure by 6 weeks of culture. However, while both 5% and 10% load constructs formed crimp, 10% load constructs formed larger, more regularly spaced crimp, indicating that there may be a dose-dependent response to load in the formation of crimp. Crimp is important to proper ligament function, however it is still unknown how it is formed. Our system could be a promising means to further explore what regulates cellular production of crimp.

As mentioned previously, crimp serves a protective role in ligaments by taking on initial loads when strains are applied [5,53,66], thus collagen crimp contributes largely to the toe region of the stress-strain curve. Here, toe region mechanics improved 2.5-3.5 fold for loaded constructs over static controls (**Fig. 5B**), suggesting the development of crimp may play a functional role in these improved toe mechanics [10]. The increase in toe region properties could also be due to increased alignment at the fibril level allowing fibrils to take on more load more quickly. However, we feel this is unlikely the case since transition strain would decrease with time in culture if increases in the toe modulus were solely due to increases in fibril alignment, and this is not the case here. However, another limitation to this study is preconditioning was not performed prior to mechanical tests which could affected the reported toe region mechanics. Preconditioning is used to normalized the load history of samples and acquire a repeatable response to load from a sample [67]. However, preconditioning can induce fibril organization in engineered tissues which can bias mechanical data [12,20,36,39,43], particularly when comparing to 0 week or unloaded constructs. Thus, tests without preconditioning are common in engineered tissues, however, future studies should use preconditioning for a better understanding of toe region properties.

Interestingly, despite both 5 and 10% load producing increased toe region properties and increased hierarchical organization, only 5% load produced a significant increase in elastic mechanics over static controls (**Fig. 5A**). As hierarchical collagen organization increases throughout development so too does tensile mechanics [11,68,69]. Similarly, in this study as hierarchical organization improved in all constructs with time in culture, so too did tensile mechanics. By 6 weeks, the elastic modulus of all groups reached the reported range of immature 1 week old bovine ACL (1-3 MPa) [46], however 5% load constructs exceeded this range (**Fig. 5A**) despite being less organized than 10% load constructs. The size and organization of collagen fibers is not the sole determining factor for tissue mechanics. It has been reported that tissue strength is reliant on collagen fiber orientation, density, diameter, and degree of crosslinking [6,70–73]. Cyclic loading, particularly 5% load significantly increased collagen content by 6 weeks of culture, with 5% load constructs reaching neonatal bovine ACL collagen concentrations (**Fig 6**). This increase in collagen content in 5% load constructs may explain the improved elastic mechanics. However, while 10% load did not increase collagen accumulation over static controls, it did stimulate increased LOX activity at 0 and 6 weeks.

LOX is an enzyme produced by cells which naturally crosslinks collagen. LOX crosslinks are another critical element in providing mechanical strength and transferring load between fibrils [5,6,70–73]. It is not well understood how mature LOX crosslinks form [74,75], but it is thought to be due to a cellular response to load [71,76,77]. In this study loading significantly increased LOX activity temporally throughout culture, with 10% load inducing a 2-fold higher activity at 0 and 6 weeks in comparison to 5% load (**Fig. 6**). It has been previously reported that cyclic loading of mature tendons stimulates or activates the cellular mechanical stretch ion channel PIEZO1 and it has been suggested this activation of PIEZO1 stimulates LOX synthesis, producing stronger tendons [77]. Further, it has been reported PIEZO1 is strain dependent in chondrocytes, with only strains higher than 8% triggering the channel [76,78–80]. While strain dependency of PIEZO1 has not been studied in the ACL, these ion channels are likely also strain-dependent in ligament fibroblasts. Thus, this may explain why 10% cyclic load in this study resulted in a 2 fold increase in LOX activity over 5% load.

Surprisingly, even though LOX activity was increased with 10% cyclic loading, there was no corresponding increase in elastic mechanics. This lack of increase in mechanics may be because LOX initially forms immature divalent crosslinks which do not have a significant effect on tissue mechanics [81,82]. With time these crosslinks can condense into mature trivalent crosslinks, which are believed to be a primary source of strength in collagen fibers [73,75,83,84]. These mature trivalent crosslinks take weeks-to-months to form *in vivo* and *in vitro* [36,74,75,82]. Therefore, the 10% load constructs may need more time in culture to form trivalent crosslinks, which may then increase tissue mechanics. One limitation of this study was we did not measure LOX crosslinks directly. Future studies should evaluate divalent, trivalent, and longer culture durations to evaluate whether increases in LOX activity lead to improved functional biochemical and mechanical properties.

Interestingly, 10% cyclic loading only significantly increased LOX activity at 0 and 6 weeks (**Fig 6**). This temporal LOX activity reflects a previous studies from our lab which found a similar release pattern of LOX activity in the media of static clamped constructs [36], and this pattern closely mirrors previously reported results in developing chick calcaneal tendons, where LOX activity increased temporally throughout development, with more dramatic increases at later stages of development [83,84]. Another explanation for this temporal LOX activity may be a change in mechanotransduction with time in culture. We hypothesize that as our constructs mature from unorganized collagen to aligned fibrils, the cells within our constructs experience the loads differently and receive higher loads [85], ultimately altering their response to the load.

In addition to differences in LOX activity with time in culture, cells differentially regulated collagen, DNA, and GAG. Collagen is the main component of ligament tissue and accumulates during development [1,5,7,57]. Intermittent cyclic load is reported to increase collagen production by cells in 2D or on aligned fibrils [23,25,41,49], but the effects at the fiber level and beyond are largely unknown. In this study, cells differentially regulated collagen production depending on degree of organization and magnitude of load, with cyclic load first producing a catabolic response early in culture when cells were in unorganized gels, which later shifted to an anabolic response once cells were anchored on aligned fibrils and fibers (**Fig. 6**). We have previously found in our system that the most collagen turnover occurs in the first 2 weeks as cells produce aligned fibrils [12,39,43]. Here, we found similar results, with loading increasing collagen turnover in the first 2 weeks resulting in reduced collagen concentrations at 2 weeks. However, once aligned fibrils were formed at 2 weeks, loading significantly increased collagen accumulation with 5% load constructs accumulating significantly more collagen than 10% load and static constructs. Previous work has reported cells produce more collagen on aligned surfaces [24], in the presences of crimp [10], and with mechanical load [10], which we also found to be the case with our cultures.

Further, it has been reported that cells sense load more intensely when attached to aligned fibers, which may lead to shifts in mechanotransduction [85]. Therefore, as cells produce aligned fibrils and fibers in our system, they may be sensing more load, which in the case of 5% cyclic load results in increased collagen production. However, there may be a threshold in cellular response. Higher loads may lead to a reduced anabolic response or a higher catabolic response [23,52], possibly accounting for the limited increase in collagen for 10% load constructs later in culture.

Further, loading differentially stimulated DNA and GAG accumulation throughout culture as well, but in an opposite manner as collagen accumulation. Intermittent cyclic loading increased DNA and GAG accumulation early in culture while cells were in unorganized collagen, but leveled off at native DNA and GAG concentration later in culture once cells were anchored on aligned fibrils and fibers, further suggesting a change in mechanotransduction. Cyclic stimulation is reported to increase cell proliferation in a dose-dependent manner with higher strains producing increased proliferation [86,87]. Here we found cyclic loading increased DNA concentrations in the first 4 weeks, suggesting increased proliferation, however we did not find a difference between 5 and 10% load (**Fig. 6**). In contrast, we did find a dose-dependent response with GAG accumulation, with 10% load significantly increasing GAG at 2 and 4 weeks over 5% and static constructs. Typically, levels above 5% strain are not used for tendon and ligament engineering because they are thought to be pathological, one sign of which is increased GAGs [88,89]. In particular, GAGs such as aggrecan and larger proteoglycans are not generally found in ligaments unless overloaded [88,89]. However, in this study, even though 10% loading produced increases in GAGs early in culture, these constructs leveled off once reaching native GAG levels, reducing the likelihood that loading induced a pathological response. Instead, this may suggest higher magnitude cyclic strains have a great effect on cell production of small leucine rich proteoglycans (SLRPs), a subset of GAGs, which have been shown to play a role in controlling fiber size and organization, and are established to increase with development and maturation [10,90]. Future studies should evaluate the type of GAGs produced to gain a better understanding of cellular regulation of SLRPs.

Here, we significantly accelerated and increased hierarchical collagen fiber formation, induced crimp formation, and increased tissue mechanics to match or exceeded that of immature ACL. Further, we have developed a system which provides an opportunity to explore how cellular mechanotransduction changes with increasing hierarchical collagen organization. However, this work is not without limitations. While these are some of the most organized ligament replacements to date, further maturation is needed to reach mature native levels. This study was an initial foray in evaluating the effect of cyclic load on cell driven hierarchical collagen organization and thus, we focused on the high and low ends of the spectrum of strain measured in native ACL (up to 5- 12%) [33]. Potentially, higher strains, longer cultures, or a combination of different loading cues are needed for further maturation. Further, the intermittent loading regime used is specifically tailored for optimal matrix turnover in unorganized gels [35,39–41], but changing any aspect such as magnitude, frequency, duration, or rest period at any point in culture could change outcomes. In particular, here we found cellular response to load varied with organization, with 10% cyclic load improving properties early in culture when cells are in unorganized gels, and 5% load driving improvements later in culture, once cells are anchored on aligned fibrils. This suggests that a load control regime or adaptive loading regimes which changes magnitude, duration, or frequency as the tissue develops may be needed to drive further maturation. However, despite these limitations, this study provides new insight into how intermittent cyclic loading affects cell-driven hierarchical fiber formation and how cells respond to these loads differentially depending on degree of maturation. A better understanding of how mechanical cues regulate fiber formation beyond the fibril level will help to develop better rehabilitation protocols to drive repair *in vivo* and improve loading protocols to drive collagen fiber maturation in engineered tissues.

## 5. Conclusions

Collectively, this study demonstrates intermittent cyclic loading at strains that reflect the native ACL environment accelerate and improve hierarchical collagen organization, crimp formation, and tissue mechanics, producing constructs that match or exceeded the properties of immature ACL. Further, we demonstrated these improvements in collagen organization and mechanics were cell driven, with the effect of cyclic load on cells varying depending on the degree of organization and magnitude of strain. We found 10% cyclic load drove early improvements in mechanics and composition when cells were in unorganized gels, while 5% load was more beneficial later in culture once cells were anchored on aligned fibers, suggesting a shift in mechanotransduction or a cellular threshold to response. Currently, it is not well understood how cells regulate hierarchical collagen fiber formation and little is known about cellular response to load beyond the fibril level [1,11,32]. This study provides new insight into how cyclic loading affects cell-driven hierarchical fiber formation. A better understanding of how mechanical cues regulate fiber formation will help to better engineer replacements and develop better rehabilitation protocols to drive repair after injury.

## Author Contribution

L.T. and J.L.P. conceived the project and wrote the manuscript. L.T. carried out experiments and analyzed the data. J.L.P. supervised the project and acquired funding. All authors edited the manuscript.

## Acknowledgements

The authors would like to thank Drs. Carl Mayer, Dmitry Pestov, and Ethan Brown for their assistance with this study. The authors acknowledge the use of facilities within the Nanomaterials Characterization Core and the Virginia Commonwealth University Cancer Mouse Models Core Laboratory, supported in part, with funding from NIH-NCI Cancer Center Support Grant P30 CA016059.

