## Supplemental for "Intermittent Cyclic Stretch of Engineered Ligaments Drives Hierarchical Collagen Fiber Maturation in a Dose- and Organizational-Dependent Manner"

**Supplemental Figures:**

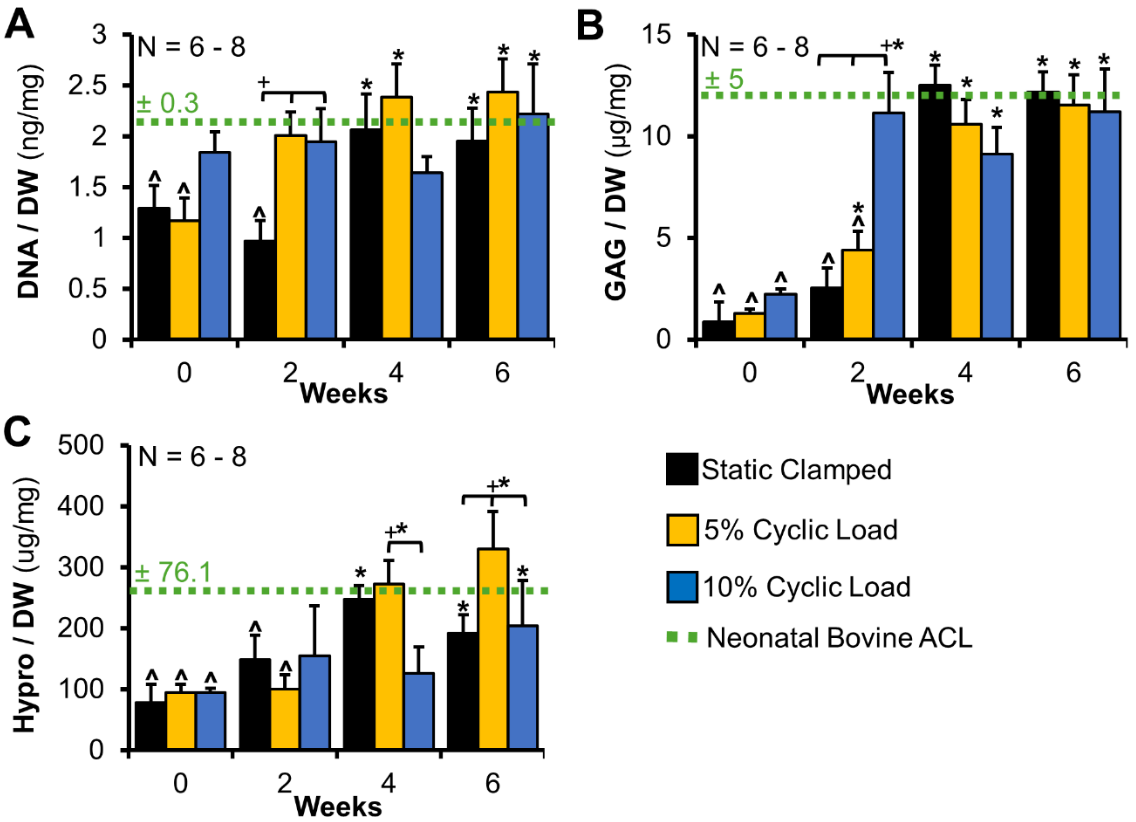


**Supplemental Figure 1**: DNA, GAG, and collagen content, represented by hydroxproline, normalized to dry weight (DW, N = 6-8). Cells appeared to have a shift in mechanotransduction with time in culture, with 5 and 10% load leading to differential changes in composition throughout culture. A-B) Loading accelerated DNA and GAG accumulation early in culture, with significant increases over static controls by 2 weeks. However, later in culture, all groups leveled off at native concentrations. C) Collagen (hydroxyproline) concentration increased with time in culture for all groups, but 5% cyclic loading produced further significant improvements by 6 weeks. Data shown as mean shown as ± S.E.M., Significance compared to *0 week, ^native, and +bracket group ($p < 0.05$)

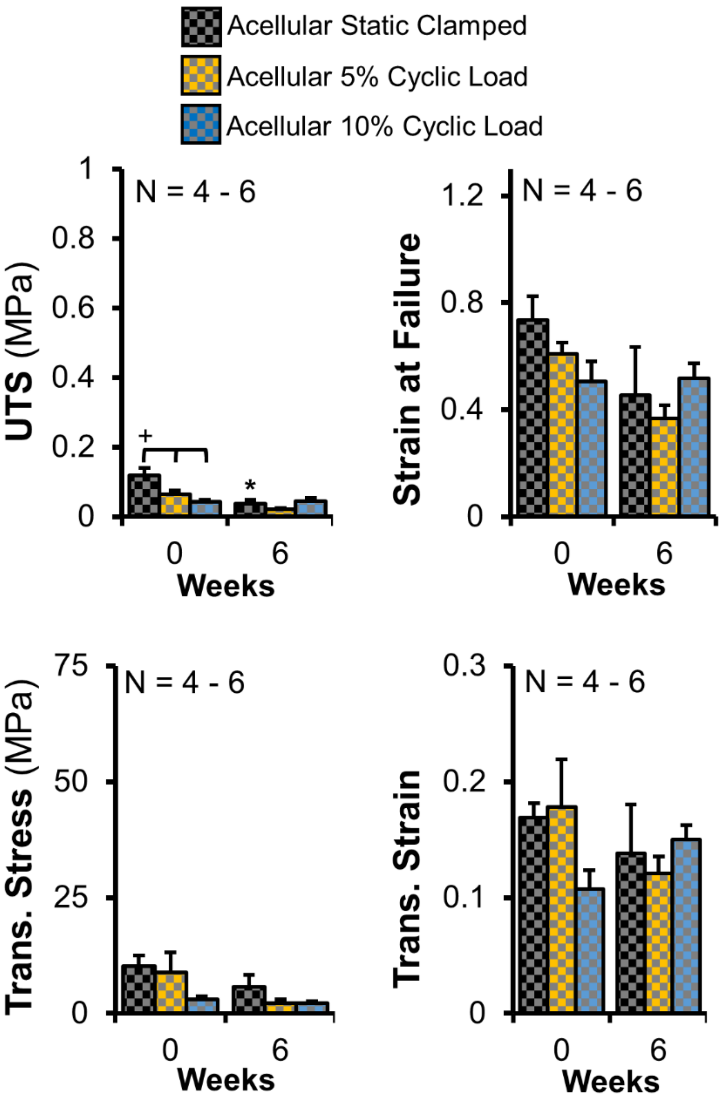


**Supplemental Figure 2**: Loading of acellular constructs drives little to no change in tensile mechanical properties, including ultimate tensile strength (UTS), strain at failure, transition stress, and transition strain. N = 4-6. Data shown as mean ± S.E.M., Significance compared to *0 week and +bracket group ($p < 0.05$)
